# LGI1 downregulation increases neuronal circuit excitability

**DOI:** 10.1101/2019.12.17.879833

**Authors:** Eleonora Lugarà, Rahul Kaushik, Marco Leite, Elodie Chabrol, Alexander Dityatev, Gabriele Lignani, Matthew C. Walker

## Abstract

LGI1 (Leucine-Rich Glioma-Inactivated 1) is a secreted trans-synaptic protein that interacts presynaptically with Kv1.1 potassium channels and ADAM23, and postsynaptically, influencing AMPA receptors through a direct link with the ADAM22 cell adhesion protein. Haploinsufficiency of LGI1 or autoantibodies directed against LGI1 are associated with human epilepsy, generating the hypothesis that a subacute reduction of LGI1 is sufficient to increase network excitability. We tested this hypothesis in *ex vivo* hippocampal slices and in neuronal cultures, by subacutely reducing LGI1 expression with shRNA. Injection of shRNA-LGI1 in the hippocampus increased dentate granule cell excitability and low frequency facilitation of mossy fibers to CA3 pyramidal cell neurotransmission. Application of the Kv1 family blocker, alpha-dendrotoxin, occluded this effect, implicating the involvement of Kv1.1. This subacute reduction of LGI1 was also sufficient to increase neuronal network activity in neuronal primary culture. These results indicate that a subacute reduction in LGI1 potentiates neuronal excitability and short-term synaptic plasticity, and increases neuronal network excitability, opening new avenues for the treatment of limbic encephalitis and temporal lobe epilepsies.

**Significant Statement:** Down-regulation of LGI1 protein increases excitability at mossy fiber to CA3 pyramidal cell synapses and in hippocampal primary cultures. These alterations alter network function and so could contribute to symptoms observed in patients affected by limbic encephalitis.

## INTRODUCTION

LGI1 (leucine-rich glioma inactivated protein 1) is a secreted brain protein and is part of the synaptic extracellular matrix (Fukata et al., 2010a). Although mutations In LGI1 were first described in association with malignant glioblastoma (Chernova et al., 1998), LGI1 mutations were later found to be highly associated with autosomal dominant lateral temporal lobe epilepsy with auditory hallucinations (ADLTE) (Kalachikov et al., 2002; Morante-Redolat et al., 2002). Mice with an embryonic knock out of LGI1 die within three weeks from birth due to severe generalised seizures (Chabrol et al., 2010b). Given its structure, with a lack of a transmembrane domain, enriched leucine repeats and cysteines at the N-terminal and a beta-propeller at the C-terminal, LGI1 is an ideal protein-protein interactor (Kobe and Kajava, 2001; Yamagata et al., 2018). LGI1 associates presynaptically with a-Disintegrin-and-Metalloprotease (ADAM) protein 23, which is part of the macro complex formed by the Kv1.1 shaker potassium channel and Kvβ1 regulatory subunit (Fukata et al., 2006; Schulte et al., 2006). It has been suggested that LGI1 regulates both the kinetics and density of Kv1.1 channels in the axon (Seagar et al., 2017). By trapping Kvβ1, LGI1 prevents the rapid closure of Kv1.1, changing its dynamics from fast opening-fast inactivation to fast opening-slow inactivation (Schulte et al., 2006). It has also been shown that in LGI1 knockout (KO) mice, the density of Kv1.1 at the axon initial segment is greatly reduced (~50%) and this affects the intrinsic properties of CA3 neurons (Seagar et al., 2017). Postsynaptically, LGI1 binds to ADAM22, which is part of the complex that links AMPA (α-amino-3-hydroxy-5-methylisoxazole-4-propionate) receptors to stargazin and PSD-95 (postsynaptic density-95) scaffolding proteins.

Specific autoantibodies against LGI1 have been also detected in the serum of adult patients with limbic encephalitis (LE), characterized by faciobrachial dystonic seizures, status epilepticus and cognitive impairments (Irani et al., 2010; Plantone et al., 2013). Plasma exchange treatment in these patients ameliorates the cognitive symptoms and the severity of the seizures, providing evidence for a causative effect of the antibodies (Crisp et al., 2016). The LGI1 autoantibody targeting protein-ligand interactions between LGI1 and ADAM22/23 has been proposed to be the basis of LGI1 autoimmune encephalitis (Ohkawa et al., 2013). Chronic infusion of IgGs-LGI1 in the ventricles of wild-type (WT) mice altered both the density of AMPA receptors and Kv1.1 channels, and was associated with behavioural deficits (Petit-Pedrol et al., 2018). However, it is unclear to what extent local inflammation rather than a direct protein interaction contributed to this.

Conditional or total KO animals for LGI1 have been previously employed to study the consequences of loss of LGI1, but these animals have developmental abnormalities which have been implicated to contribute to the phenotype (Chabrol et al., 2010b; Fukata et al., 2010b; Yu et al., 2010; Boillot et al., 2014). In this study, we asked if a subacute local reduction in LGI1 was sufficient to increase network excitability. First, we used an RNA interference (shRNA) approach to silence LGI1 by injecting lentiviral particles expressing the shRNA against LGI1 in vivo to target the mossy fiber – CA3 synapses, where there is a high expression of Kv1.1. There we examined short-term plasticity and neuronal excitability. We then investigated LGI1 shRNA effects on neuronal circuits by using a combination of calcium imaging and multielectrode arrays (MEAs) *in vitro*. Collectively, our results indicate a central role for LGI1 in regulating network excitability and provide a link between deficiency in LGI1 and the emergence of seizure activity.

## Materials and Methods

### Animals

Animal care and procedures were carried out in line with the UK Animals (Scientific Procedures) Act, 1986 and under the Home Office License PLL/PIL 11035/13691. Animals were kept at room temperature and maintained on a 12/12 hours’ dark/light cycle with free access to food and water. Experiments were carried out after a minimum period of 7 days of habituation.

### cDNA cloning

All the experiments were performed using a pLentivirus_mU6_shRNA-LGI1_PGK_EGFP or a pLentivirus_mU6_scramble_PGK-EGFP (see supplementary table 1 for sequences). Three shRNA plasmids to knockdown mouse *LGI1* (GeneID:56839) were generated by the insertion of three different siRNA sequences (obtained from Dharmacon, Horizon Discovery, supplementary table 1) into AAV U6 GFP (CELL BIOLABS INC., San Diego, CA 92126, USA) using BamH1 and EcoR1 restriction sites. Positive clones were confirmed by sequencing. Furthermore, mouse neuroblastoma cell line Neuro2a (N2a) were transfected with the shRNA plasmids along with a control plasmid using calcium phosphate method (Kwon and Firestein, 2013) to investigate the knockdown efficiency of the constructs. After 48 hours of transfection, cells were harvested to isolate the RNA using EURx_Genematrix kit (ROBOKLON GMBH, Berlin, Germany) as per manufacturer’s protocol. RNA was further converted to cDNA using High Capacity cDNA Reverse Transcription Kit (Applied Biosystems, now ThermoFisher Scientific) and used to quantify the mRNA levels of LGI1 using RT-qPCR. GAPDH gene was used for normalization. The original plasmids were then subcloned into a pCCL_Flexed_MCS backbone using *SalI* and *EcoRI* restriction sites. The high titer and ultra-purified viruses were produced by VectorBuilder (LGI1-shRNA titer 3.08 × 10^9^ IU/ml; Scr-shRNA titer 4.25 × 10^9^ IU/ml). The commercial viruses were expressed in primary cultures with a transduction rate >50% estimated by counting the EGFP positive neurons relatively to the total number of DAPI positive cells.

### Neuronal cultures and transduction

Mixed neuronal and glial cell cultures were prepared from cortical and hippocampal embryonic (E18) C57Bl/6J mice (Envigo) as previously described (Dixon et al., 2015). Briefly, the embryos were extracted from the amniotic sac of the pregnant mouse, sacrificed and the brain was placed in an ice-cold Hank’s balanced salt solution (HBSS, Thermo Fisher, Invitrogen). Meninges were then removed, cortex and hippocampi were isolated, trypsinized for 3 minutes at 37°C and mechanically disrupted. Neurons were then plated on glass coverslips (WVR international) previously coated with PLL (Poly-L-lysine hydrobromide, mol wt 70.000-150.000, Sigma Aldrich) diluted to 1mg/ml in borate buffer (0.1M boric acid, pH 8.5) at 400.000 cells/coverslips. Cells were initially plated in DMEM-10% FBS (Thermo Fisher) for two hours in a humidified incubator containing 5% CO_2_ and 95% oxygen. Complete Neurobasal medium containing 1% PenStrep (penicillin 10.000 U/mL - streptomycin 10mg/mL, Sigma Aldrich), 1% GlutaMAX™ (Thermo Fisher) and 2% B27 supplement (Gibco) was then replaced to fill 2/3 of the well. At DIV1, cells were transduced with the viruses using 5 MOI (multiplicity of infection) and checked a week later for detection of the EGFP expression. Half media was replaced with a fresh one every 7 days at 37°C and neurons were used at DIV 17-23.

### Western blot

Samples were lysed in RIPA buffer (150 mM sodium chloride, 1.0% Triton X-100, 0.5% sodium deoxycholate, 0.1% sodium dodecyl sulfate, 50 mM Tris, pH 8.0; Sigma Aldrich) completed with a protease inhibitors cocktail (Roche Diagnostics GmbH). Prior to loading, samples were boiled at 95°C for 5 minutes, diluted in 5M Urea (Sigma) solution and 20μg of protein were loaded in each lane. Samples were loaded into premade gels (10% Bolt Bis-Tris Plus Gels, Invitrogen). Proteins were transferred on to Nitrocellulose membrane using a semi-dry method (Chabrol et al., 2010b). Nitrocellulose membranes were washed in PBS (phosphate-buffered saline) incubated with primary antibodies and secondary HRP antibodies in PBS-T (0.1% Tween20) containing 1% milk. The following antibodies and dilutions were used: 1:250 polyclonal goat-anti-LGI1 (Santa Cruz, c-19), 1:4000 monoclonal mouse-anti β-actin (Sigma), 1:1000 rabbit-anti-GFP (Abcam ab6556). Chemiluminescence signals were revealed using PierceM ECL substrate (Thermo Fisher). Images were acquired using a ChemiDoc™ Imaging System (BioRad) and analysed with Image Lab software. LGI1protein levels were normalized over β-actin from the same sample lane.

### Immunofluorescence

Primary hippocampal neurons were fixed for 10 minutes in 4% paraformaldehyde, rinsed in PBS 3 times for 5 minutes, permeabilized in 0.1% Triton X-100 (Thermo Fisher) for 10 minutes and blocked with 5% BSA (Sigma Aldrich) for 30 minutes. Primary antibodies were applied overnight using guinea pig vGLUT1 1:1000 (Millipore, ab5905) and rabbit Homer1 1:500 (Synaptic System, 160 033). Hoechst 33342 was used as nuclear stain and then coverslips were mounted on a droplet of mounting media (Sigma Aldrich) on a glass microslide (VWR). Images were acquired using a 20 × objective at a Zeiss confocal microscope.

### Live cell imaging: viability assay

Propidium iodide (PI) dye (1ug/ml, Thermo Fisher) and Hoechst 33342 dye (4.5uM, Thermo Fisher) were incubated in the cell media for 20 minutes in the incubator. Coverslips were then moved into a metallic chamber and imaged at with a Zeiss confocal microscope using a 20X objective. Images were then analysed by an automatic ROI detection plugin in ImageJ software (Rueden et al., 2017).

### Live cell imaging: calcium imaging

Calcium imaging experiments were performed with mouse embryonic (E18) cortical cultures between 21 and 23 DIV. Cells were initially incubated for 30 minutes at room temperature with 5 μM Fura-2-AM and 0.005% pluronic acid (ThermoFisher) in HEPES-aCSF solution (125 mM NaCl, 2.5 mM KCl, 2 mM MgCl2, 1.25 mM KH2PO4, 2 mM CaCl2, 30 mM glucose and 25 mM HEPES, pH 7.4). Cells were recorded for 10 minutes using a 20x inverted objective (Olympus IX71). Excitation light was provided by a xenon arc lamp, the beam passing through a monochromator at 340 and 380 nm with a bandwidth of 10 nm (Cairn Research, Faversham, UK). Emitted fluorescent light was detected by a cooled CCD camera (QImaging Retiga EXi, 1394) through a 510 nm long-pass filter (515 nm dichroic). Images were initially processed using Andor software (Belfast, UK) and analyzed with a custom Python script. The script cycles through all the individual cells and computes the baseline as the 5th per centile of the whole data set array. Each peak was detected as a local maximum of 20% greater than the average baseline. Active cells were counted as those cells which have at least 1 peak over 60 frames. The frequency of peaks was computed as a total number of peaks divided by the data size (60 frames).

### Multielectrode Arrays

A single MEA device consisted of a 6-wells multielectrode array formed by 60-channel (54 recording electrodes plus 6 internal references) (MultiChannel Systems, MCS, Reutlingen, Germany). The MEA was heated up at 37°C for all the time of the experiment. The signal was digitized at 25 kHz. A parafilm cap was made in order to avoid evaporation and changes in the osmolality of the media (Blau *et al.*, 2009). The bath was grounded with a silver chloride wire wrapped around one of the name-pin of the MEA device and placed in well A for all the experiments. MEAs were sterilised by the use of 1% solution of Terg-a-zyme (Sigma Aldrich). MEAs coating was done by placing a mixture of laminin (0.05 mg/ml, ThermoFisher) and PLL (30.000-70.000 MW, Sigma Aldrich) at ratio 1:2 in borate buffer at the center of the device (Colombi *et al.*, 2013). Cells were plated at 45.000 neurons per well. Neurons were transduced at DIV1 and the 6 wells were equally divided by treatments. Recording protocol was organised in 5-minute recovery followed 10 minutes recording. Collected data were processed using the Spycode software (Matlab) kindly shared by Ilaria Colombi (Dr. Michela Chiappalone’s lab, Italian Institute of Technology, Italy). The results of analysis were statistically analysed and presented using GraphPad Prism7. Ten standard deviations of the noise were defined as the threshold to detect single extracellular spikes (Colombi et al., 2013).

### Solutions (for whole-cell clamp and LFP experiments)

The cutting solution used for LFP and whole-cell experiments to perfuse the animal while cutting the brain contained (mM): sucrose (105), NaCl (60), NaHCO_3_ (26), glucose (15), NaH_2_PO_4_ (1.25), KCl (2.5), ascorbic acid (1.3), sodium pyruvate (3), CaCl_2_ (0.5), MgCl_2_ (7). Ringer solution was used for LFP experiment as slice storage and recordings in LFP and whole-cell experiments (mM): NaCl (125), NHCO_3_ (26), glucose (15), NaH_2_P0_4_ (1.25), KCl (2.5), MgSO_4_ 7 H_2_O (1.3), CaCl_2_ (2). Both solutions were bubbled with 95% O_2_/5% CO_2_ to have 7.4 pH and osmolality of 305-310 mOsm. Whole patch-clamp internal solution contained (mM): K-gluconate (K-Glu, 135), KCl (4), Hepes (10), Mg-ATP (4), Na-GTP (0.3), Na_2-_ phosphocreatine (10). The pH was 7.3 and osmolarity was 291-295 mOsm.

### Electrophysiology: whole cell patch-clamp and local field potential

Five-six weeks old male mice were deeply anesthetised with isofluorane (3% in oxygen, 2 L/min) injected with a sub-lethal dose of pentobarbital IP (140 mg/kg) and then perfused with ice-cold sucrose solution. The brain was rapidly removed and placed in a glass petri dish filled with bubbled ice-cold sucrose solution in order to extract the intact hippocampi, which were then embedded in a 3% agar support, angled at 30° with respect to the dorso-ventral axis, and sliced with a Leica VT1200 vibrating blade microtome. Isolated hippocampal transverse slices (400 μm) were then transferred to a submerged chamber (95% O_2_, 5% CO_2_) placed into a warm bath (34°C) for 20 minutes and then placed at room temperature for 60 min in standard recording aCSF before experiments started. Slices were submerged into the recoding chamber of an upright Olympus BX50WI microscope. Slices were continuously perfused and bubbled in extracellular recording solution (23-25°C) and visualised with 10X and 40X objectives (Olympus) using DIC optics. The recordings were performed with borosilicate thin glass electrodes (4-6 MΩ, vertical puller Narishige PC-10), filled with filtered cold K-gluconate solution and connected to an Axon Multiclamp 700B amplifier (Molecular Devices). The signal was low-pass filtered at 2kHz (Bessel) and sampled at 10kHz using WinEDR (John Dempster, University of Strathclyde). Cells with RMP (resting membrane potential) > −55mV, access resistance > 20 MΩ, bridge balance > 10 MΩ and leak current > 200 pA were discarded. Recordings were not corrected for liquid junction potentials. Neurons were maintained at −70mV for the whole time of the recording. Traces were analysed with a bespoke custom-made Python script. Neuronal excitability was tested in current-clamp by injection of increasing current steps (100ms initial delay, 1000ms pulse with 10pA step increase from −10pA up to 100pA).

For LFP experiments. slices were initially left to equilibrate for 1 hour. Stimulating bipolar electrodes were placed on the granule cell layer of the DG, meanwhile, a recording field electrode filled with aCSF was placed in CA3 stratum lucidum. Square-wave 80 μs-long pulses were delivered using a constant current stimulus isolator (DS3, Digitmer, Hertfordshire, UK). The recording electrode was made of borosilicate glass (1-3 MΩ) and filled with an extracellular perfusion solution. The position of the stimulation electrode was adjusted to maximize the fEPSP amplitude and stimulation intensity was adjusted to elicit fEPSPs that were 1/3 of the maximum amplitude. Slices were discarded if it was not possible to elicit a fEPSP with an amplitude of at least 1mV. Paired pulses were delivered with 50 ms stimulus interval. The short-term plasticity protocol chosen for assessing the MF-CA3 circuit, consisted of 10 repeated stimuli at 0.05 Hz, followed by 40 stimuli at 1Hz (Chandler *et al.*, 2003). This “wind-up” stimulation paradigm was elicited twice in a row and signals were averaged during off-line analysis. Wind-up ratio was obtained by normalizing the facilitation by the average baseline amplitude. After the wind-up protocol, in all experiments, the pathway was confirmed as the mossy fibre pathway through the application of the group II metabotropic glutamate receptor agonist, 1μM DGC-IV (Tocris Bioscience). This was then followed by, the AMPA/kainate receptor antagonist 50μM NBQX (Tocris Bioscience) (Chandler *et al.*, 2003) in order to isolate the fiber volley signal which was then subtracted off-line from fEPSP recordings with a custom script in Python 3.0 and slopes were analysed off-line. 1μM α-Dendrotoxin (Alomone Lab) was dissolved in 0.1% BSA and acutely applied to the slice. The signal was low-pass filtered at 2kHz and sampled at 20kHz using WinWCP (John Dempster, University of Strathclyde).

### In vivo stereotaxic injection of substances

C57BL/6J WT male mice (4 weeks old, 16-18 gr) were anesthetised with isoflurane (3% in 2 L/min O_2_), and injected with Metacam (0.13 mg/kg) and Buprenorphine (0.02 mg/kg, 40 minute since onset) (Temgesic, Scherig-Plough, UK) subcutaneously. Two burr holes were drilled (WPI) according to the coordinates from bregma suture (mm): (AP −2.3, ML ±3, DV1 −2.2, DV2-2.8) to target the central hippocampus. The coordinates were adjusted in proportion to the distance between the bregma and lambda sutures over a max ML distance of 4mm. One μl of each virus were injected at 100nL/min. After infusion, the micropipette remained in place for 5 minutes to avoid backflow of the virus to the surface. Then the mouse was monitored until fully awake (Counsell *et al.*, 2018).

### Statistical analysis

All the data are expressed as mean values ± SEM (standard error of the mean), if not stated otherwise. The normality of data distribution was assessed using the Kolmogorov–Smirnov normality test. For normally distributed data, Student’s unpaired t-test or paired t-test were used, as appropriate. Analysis of the variance (ANOVA)’s test coupled with post-hoc correction, was used when 3 or more normally distributed data were compared. Data not normally distributed were compared using the Mann-Whitney test. Range and IQR (inter quartile range) were used to indicate as a measure of the dispersion of the data around the median (Norman and Streiner, 2000). The data are represented by box and whiskers graphs: bars symbolised minimum and maximum value, box edges were the 25^th^ and 75^th^ percentiles, the horizontal central line represents the median, the “+” symbol represents the mean. Statistical analysis was done by employing GraphPad© Prism 6 software, Microsoft Excel 2010 (Windows), Matlab © (The Mathworks, Natick, MA, USA), Clampfit Axon pClamp™ and Python 3.1© (open-source software available at www.python.org/) with Anaconda© AI/ML enablement platform. All the experiments were performed blinded for the treatment group.

## Results

### Downregulation of LGI1 increases mossy fiber facilitation

We used an shRNA approach to subacutely decrease LGI1 expression, over the course of 7 days. To optimise the knock down (KD) sequence, we initially evaluated three shRNA-LGI1 sequences in mouse neuroblastoma cell line (Neuro2a). All three sequences resulted in ~80% KD efficiency compared with the effect of the scramble sequence (Supplementary Figure 1). For further experiments, we used shRNA#2 cloned in a lentiviral vector driven by a U6 promoter, (LGI1-shRNA; Figure 1A). We transduced primary neuronal cultures at DIV1 with both LGI1-shRNA and Scr-shRNA, and collected them at DIV21 for quantification of LGI1 protein. LGI1 levels were reduced on average by 38.2 ± 0.05% in LGI1-shRNA cultures compared to Scr-shRNA ones (Scr-shRNA mean = 0.955 ± 0.084; LGI1-shRNA mean = 0.573 ± 0.057; p=0.05 two-tailed Student’s t-test) (Figure 1B).

**Figure 1.**
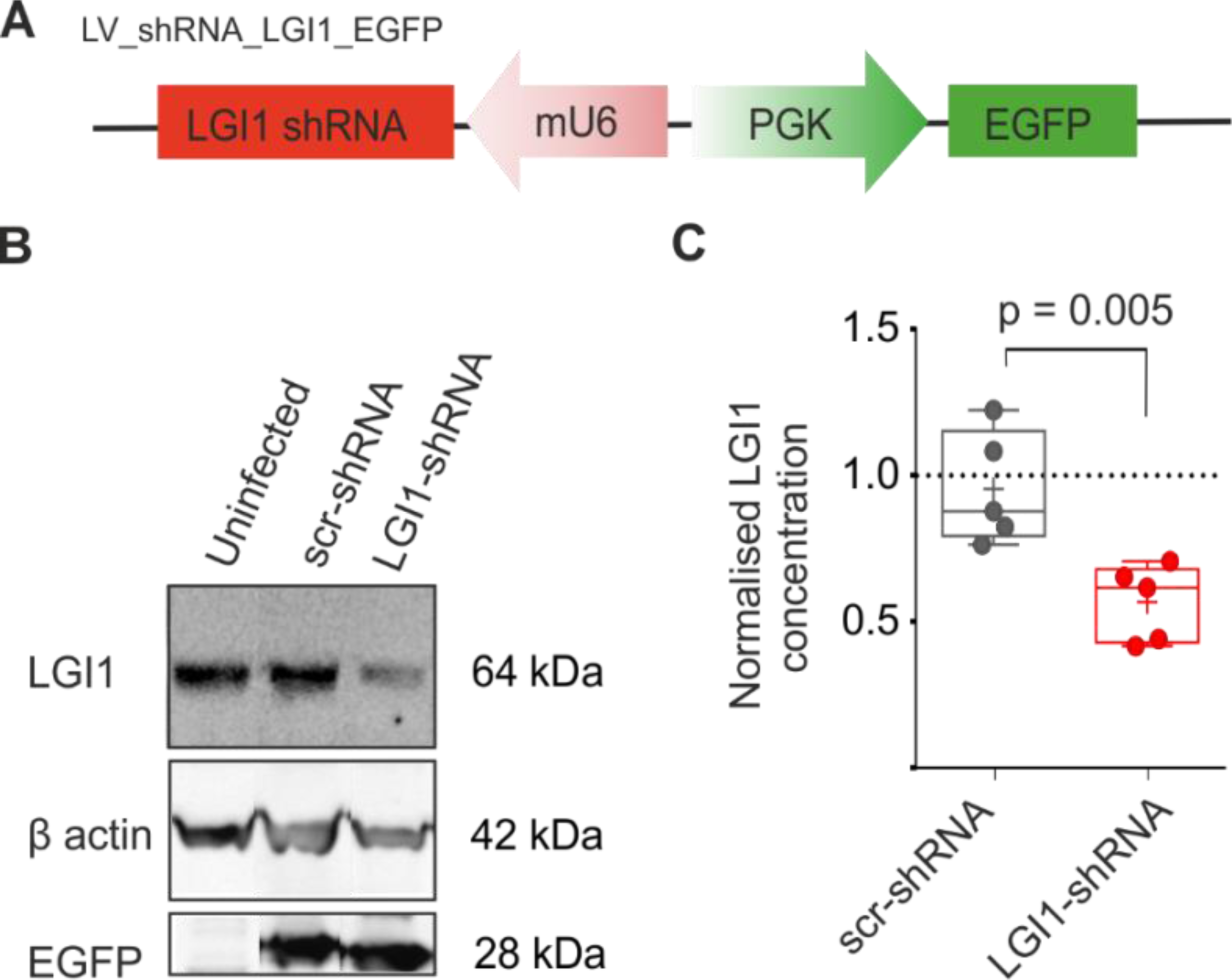
ShRNA plasmid design and in vitro LGI1-shRNA downregulation. A) shRNA construct: LGI1-shRNA is under the promoter mU6 (murine U6 small-non-coding-RNA), meanwhile the EGFP (enhanced green fluorescent protein) is under the independent promoter PGK (phosphoglycerate kinase). NeoR/KanR (Neomycin/Kanamycin Resistance). **B)** LGI1 expression in neuronal cultures. The first lane is the endogenous level of LGI1 in untreated primary cultures, the second lane shows the levels of LGI1 in cultures transduced with scramble virus and the fourth and last lane shows the levels of LGI1 in cultures transduced with shRNA-LGI1. The β-actin and EGFP signals are also shown. EGFP intensity was detected to evaluate that the reporter gene was also expressed at the same time as the shRNA and hence could be a reliable source of transduction rate. **C)** Quantification of LGI1 concentrations detected by the western blot method. p=0.005 two-tailed Student’s t-test (blots n=5 from 5 distinct preparations).

We next asked if the downregulation of LGI1 is sufficient to alter hippocampal excitability. Previous experiments on LGI1 KO have demonstrated a developmental role for LGI1 but do not directly address the impact of a subacute reduction. We tested the impact of LGI1 knockdown on dentate granule cell mossy fiber bouton to CA3 pyramidal cell synapses, as dentate granule cells strongly express LGI1 protein, suggesting a key function of this protein in those circuits (Schulte et al., 2006).

We, therefore, injected either Scr-shRNA or LGI1-shRNA in the DG region and performed local field potential experiments stimulating granule cell layers and recording in stratum lucidum of CA3. We used an established protocol (wind-up) to induce short-term MF plasticity (Chandler et al., 2003). The results show that mossy fibers in LGI1-shRNA injected hippocampi facilitated to a significantly greater degree than did Scr-shRNA injected hippocampi (Figure 2 C, D) (Scr-shRNA mean = 1.89 ± 0.04; LGI1-shRNA = 3.17 ± 0.04, p < 0.0001 two-tailed Student’s t-test). At the end of every stimulation, the specific metabotropic glutamate receptor 2 agonist, DCG-IV was applied to check that the field potential recorded was mediated by mossy fibers (Kamiya et al., 1996; Chandler et al., 2003). Scr-shRNA and LGI1-shRNA were blocked to similar degrees (Scr-shRNA mean = 0.05 ± 0.01% (~95%); LGI1-shRNA mean 0.03 ± 0.01% (~98%), p > 0.4 two-tailed Student’s t-test) (Figure 2 E). As a further control, we assessed that there was no difference in baselines slopes before and after the wind-up in control conditions (Scr-shRNA baseline#1: 0.032 ± 0.008, Scr-shRNA baseline#2: 0.039 ± 0.015, p>0.05 two-tailed Wilcoxon test).

**Figure 2.**
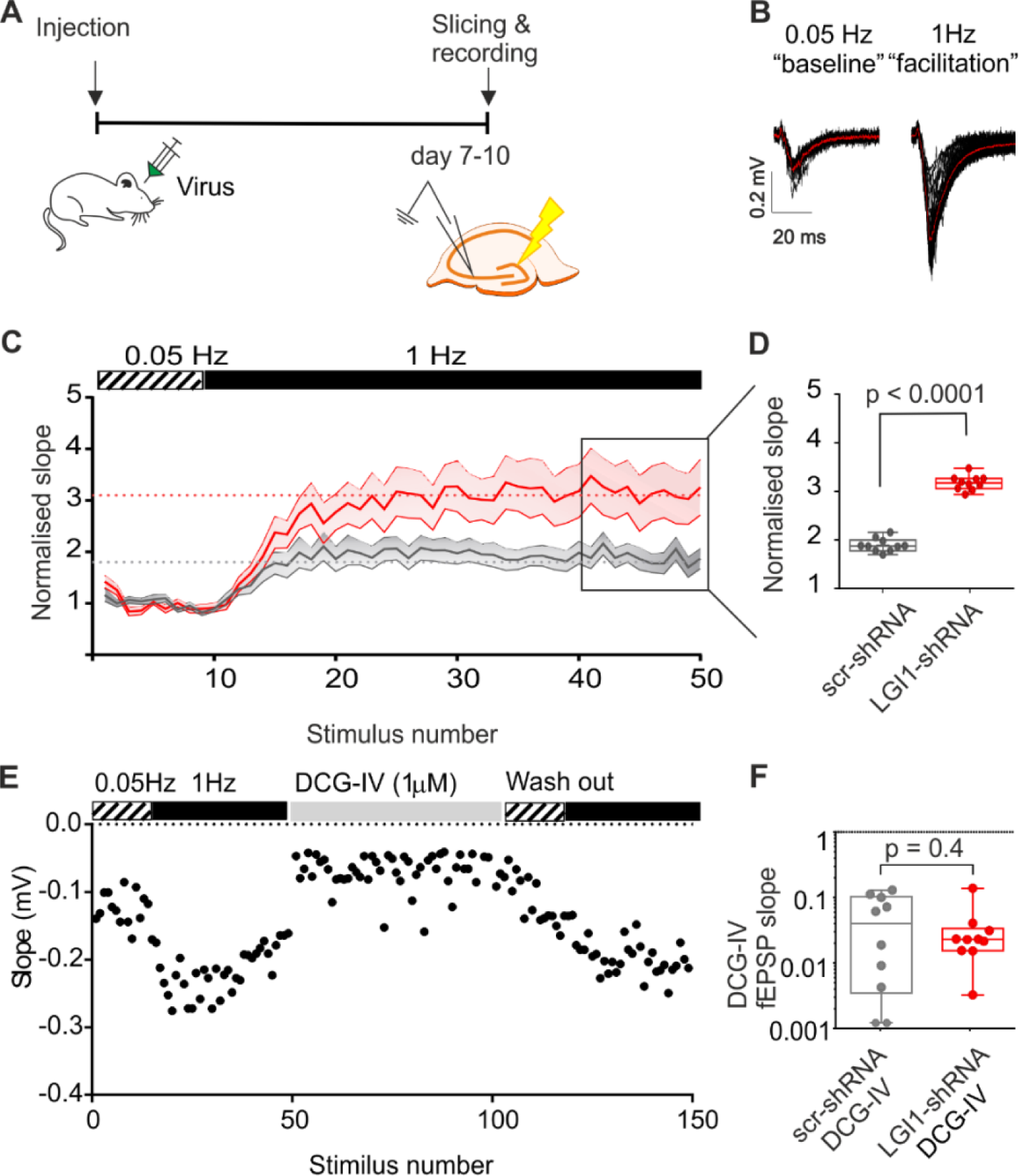
MF to CA3 local field potential is increased in animals knocked down by shRNA-LGI1. **A)** Experimental timeline and **B)** stimulation paradigm. On the right side are shown original representative trace of baseline and wind-up stimulation of MF-CA3 circuit. Red trace represents the averaged signal of 10 baselines traces and 40 facilitation traces. Black lines represent single recordings. **C)** Pooled time-line data showing the increase in slope following different stimulation frequencies. In 1Hz stimulation, the amplitude in LGI1 conditions is 30% greater than in the scramble conditions. The bar graph represents the averaged last 10 traces of each recording. (**D**) Scr-shRNA n= *27 slices from 8 animals* and LGI1-shRNA *n = 20 slices from 9 animals* **E)** representative recording showing the effect of 1 µM DCG IV in reducing fEPSP slope. (**F)** Bar graph summarising the effects of 1 µM DCG-IV in Scr-shRNA and LGI1-shRNA slices.

Lastly, short-term plasticity was also assessed by paired-pulse stimulation. This form of rapid plasticity in the DG is generally reported as facilitating and it is dependent upon presynaptic mechanisms such as calcium accumulation after the first stimulus, but also potassium channel activity (Meneses et al., 2016). Kv1 channels, indeed, are a major player in the presynaptic modulation of mossy fiber synapses (Bialowas et al., 2015; Rama et al., 2015; Meneses et al., 2016). The results show that there was no significant difference in paired-pulse ratio in LGI1-shRNA and Scr-shRNA at 50 ms delay (Scr-shRNA median = 1.99, range = (0.86 to 8.33) 7.47, IQR = 2.33; LGI1-shRNA median = 2.00, range = (0.85 to 4.66) 3.81, IQR = 1.46, p > 0.5 Mann-Whitney test. (Supplementary Figure 2).

### Blocking Kv1.1 potassium channels occludes increased facilitation due to LGI1 downregulation

Recent work has demonstrated that LGI1 expression can determine the amount of Kv1 channels at the synapse and axon initial segment, and that heterozygous mice for LGI1 (+/-) expressed 40% fewer Kv1.1 compared to that in control mice (Seagar et al., 2017). Previous work has reported that non-secreted LGI1 proteins are also able to change Kv1.1 kinetics through direct block of the Kvβ1 regulatory subunit (Schulte et al., 2006). The Kv1 family and LGI1 protein largely overlap in their expression in the medial perforant path, granule cell molecular layer and axons and terminals of DG mossy fibers (Trimmer and Rhodes, 2004; Schulte et al., 2006; Trimmer, 2015). To test if the increased facilitation was due to changes in the activity of the Kv1.1 channel, we repeated the wind-up experiments applying a broad blocker of the Kv1 family, α-Dendrotoxin, (Geiger and Jonas, 2000). In the Scr-shRNA condition, application of DTX increased the wind-up facilitation (Figure 3 A-D), while, in LGI1-shRNA condition, it decreased facilitation (Figure 3 A-B, E-F) (Scr-shRNA α-DTX mean = 32.8± 5.3% larger than with aCSF; LGI1-shRNA α-DTX mean = 67.8 ± 6.2% smaller than with aCSF; p < 0.0001, paired Student’s t-test. Scr-shRNA n = 14 from 6 mice, LGI1-shRNA n= 11 from 6 mice). As a result, there was no significant difference between wind-up facilitation in Scr-shRNA and LGI1-shRNA treated hippocampi in the presence of dendrotoxin (p = 0.7). As a control, 0.1% BSA alone had no effect on the rate of facilitation observed in two slices (WT + 0.1% BSA = 7.85 ± 0.08%, n=2 slices from 1 mouse, data not shown).

**Figure 3.**
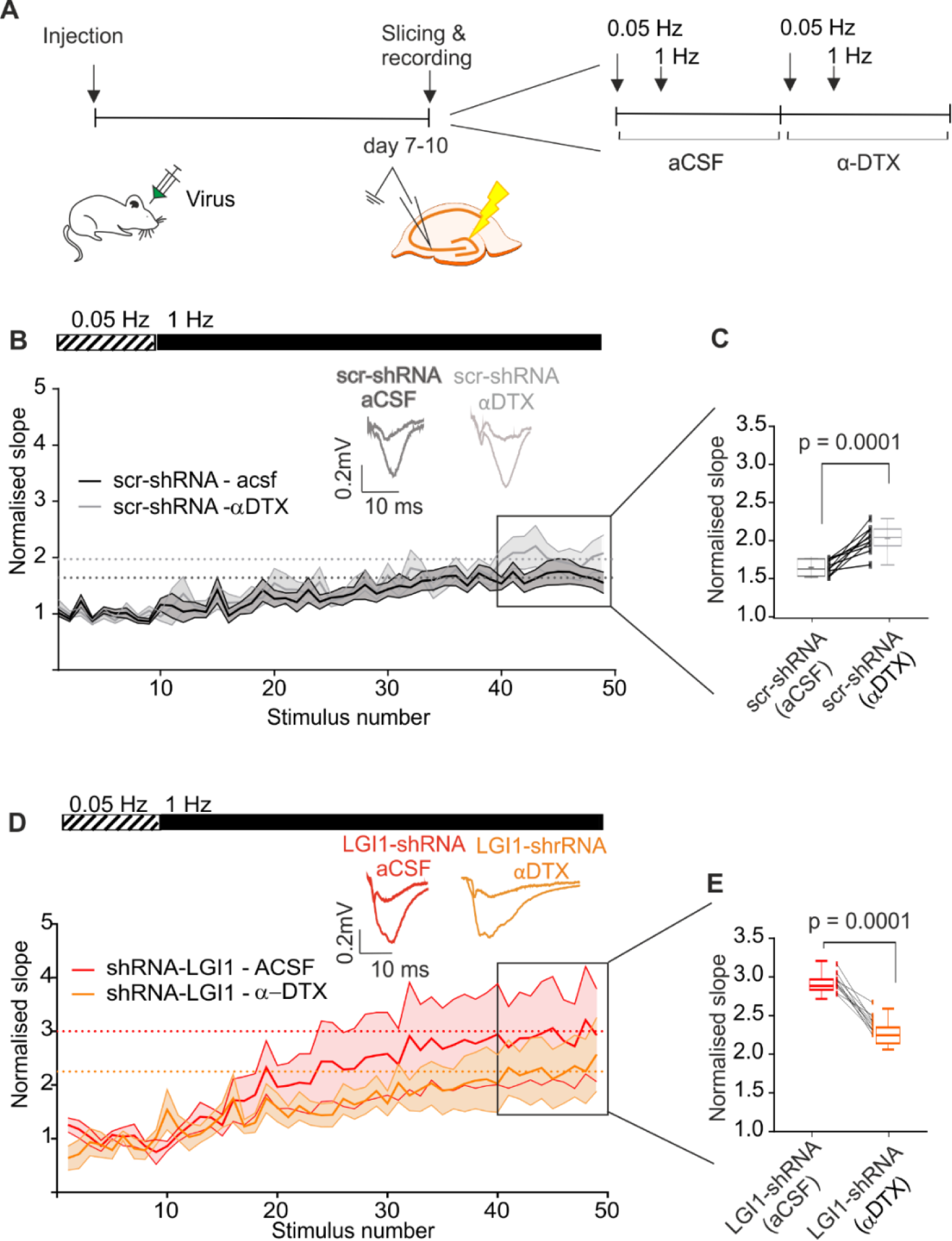
The over facilitation observed in animals with subacute reduction of LGI1 in MF to CA3 circuits is due to early inactivation of Kv1.1 channels. **A)** Experimental timeline and paradigm. Representative trace and pooled time line data of wind-up stimulation of MF-CA3 circuit before and after application of α-DTX in scramble-shRNA injected slices **(B,C)** and in shRNA-LGI1 injected slices **(D,E)**. Bar graphs in C &E show the quantification of the slope of the last 10 wind-up stimuli normalised over baseline stimulation. Scr-shRNA: n = 14 from 6 mice, LGI1-shRNA: n= 11 from 6 mice.

α-DTX had no significant effect on baseline fEPSP (baseline slope ACSF vs. α-DTX, repeated measures two-way ANOVA, p = 0.1, F (1, 20) = 1.8). Nevertheless, LGI1-shRNA showed a trend in having a smaller baseline fEPSP compared to control (Scr-shRNA n= 13 from 6 mice, LV-shRNA n= 9 from 6 mice) (Supplementary figure 3 A). The analysis of the fiber volleys showed that for both groups, after application of α-DTX the slope of the fiber volley was reduced by 12.7 % of the baseline for Scr-shRNA and 12.6 % for LGI1-shRNA (Scr-shRNA n= 12 from 5 mice, LV-shRNA n= 7 from 5 mice), but this was not significantly different (fiber volleys slope ACSF vs. α-DTX, repeated measures two-way ANOVA, p = 0.18, F (1, 17) = 1.9) (Supplementary figure 3 B).

We have therefore shown that blocking the Kv1 family occludes the increased MF facilitation in LGI1-shRNA hippocampi while having a non-significant effect on baseline neurotransmission.

### Paracrine effects of subacute LGI1 downregulation in DG granule cells

Since restricted downregulation of LGI1 is able to increase circuit excitability despite the low efficacy of LV transduction, we asked if a paracrine effect could exist in the DG. Previous work has demonstrated that LGI1 is a secreted protein, probably released from the dendritic and axonal compartment of neurons (Lovero et al., 2015). The total knockout of LGI1 from the early embryonic stage has many different effects on neuronal physiological properties, spanning both pre- and post-synaptic features (Boillot et al., 2016; Fukata et al., 2010; Zhou et al., 2009). Interestingly, when LGI1 is reintroduced onto LGI1 KO neurons, the AMPA/NMDA ratio was rescued in both transfected positive neurons and in nearby cells as well (Lovero et al., 2015). Since it is well established that excitatory neurons deprived of LGI1 have a lower rheobase and increased excitability both in KO animals, mutants and conditional KO (CAMKIIα-LGI1 cKO), we aimed to test whether subacute downregulation of LGI1 had a paracrine effect on active and passive neuronal excitability properties in neighbouring DG granule cells (Zhou et al., 2009; Fukata et al., 2010a; Boillot et al., 2014; Seagar et al., 2017). We assessed neuronal excitability of cells that were in close proximity to EGFP positive neurons (Figure 4 A-C). Neighbouring LGI1-shRNA positive DG neurons displayed a lower threshold to elicit the first AP, compared to neighbouring Scr-shRNA neurons (Scr-shRNA vs. LGI1-shRNA: p = 0.001, repeated measures two-way ANOVA with Bonferroni correction for comparisons F (1, 25) = 12.62). The half-width of the action potential measured at current threshold was not significantly different between the two groups (Scr-shRNA mean = 1.64 ± 0.58 ms; LGI1-shRNA = 1.67 ± 0.66 ms; p = 0.739, two-tailed Student’s t-test) (Figure 4 D), the latency to the first spike between the first 4 spikes over 90pA injected current was shorter for LGI1-shRNA neighbouring neurons (Scr-shRNA vs. LGI1-shRNA: p < 0.0001, repeated measures two-way ANOVA with Bonferroni correction for comparisons F (3, 57) = 25.88) (Figure 4 E). Importantly, analysis of passive properties of patched neurons highlighted a significant difference in the input resistance (Scr-shRNA vs. LGI1-shRNA: p = 0.002, two-tailed Student’s t-test), leaving the membrane capacitance unaffected (Scr-shRNA mean = 1.158 ± 0.028 pF; LGI1-shRNA = 1.244 ± 0.03 pF; p = 0.06, two-tailed Student’s t-test) (Figure 4 F, G). Resting membrane potential was not significantly different between treatments (Scr-shRNA = −66.4 ± 2.4 mV, LGI1-shRNA = −70.6 ± 2.5 mV; p = 0.25 two-tailed Student’s t-test).

**Figure 4.**
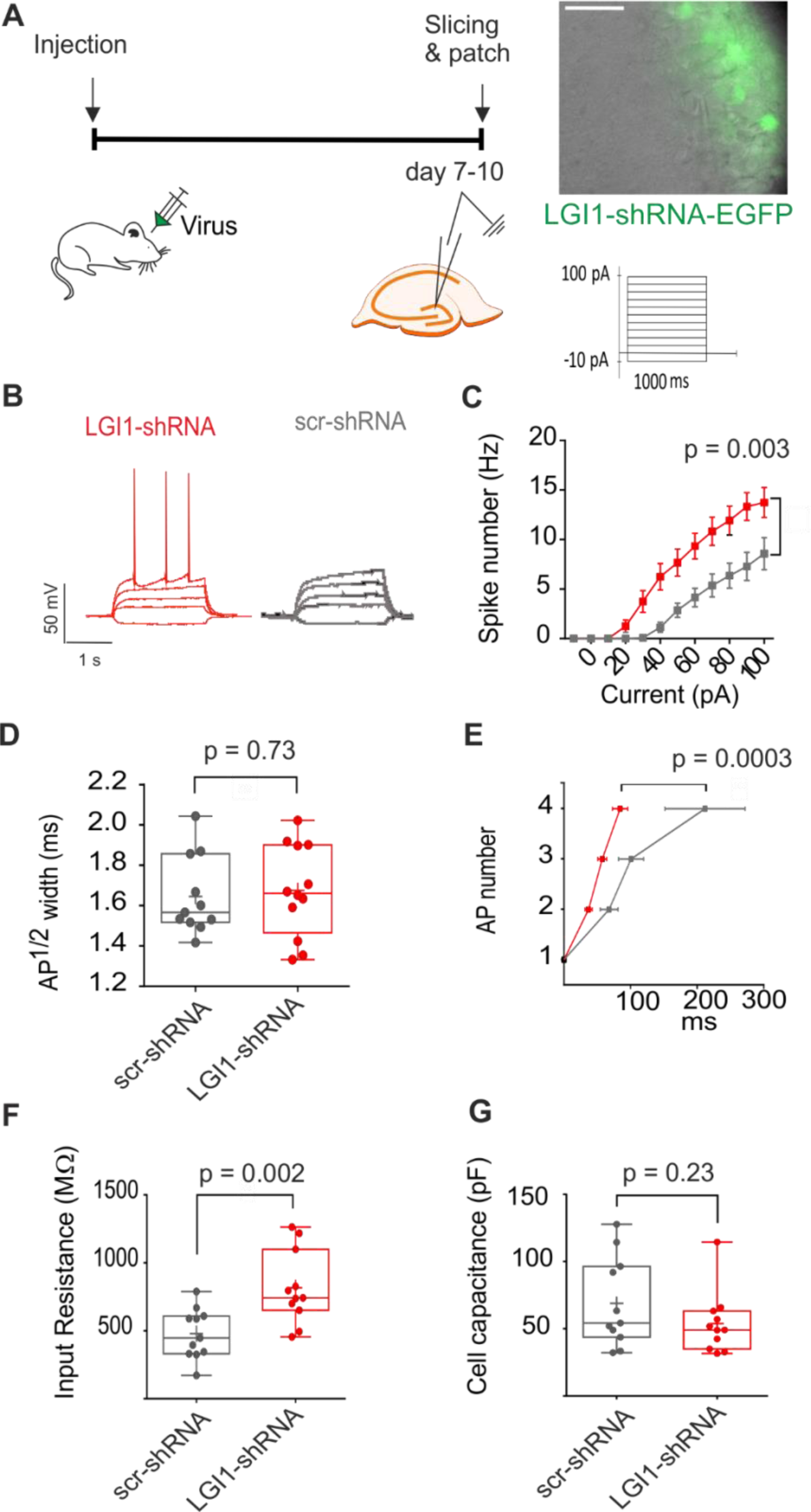
Subacute knock down of LGI1 increases intrinsic neuronal excitability. **A)** Experimental timeline and paradigm. The image shows granule cells in an acute slice. The pipette is patching an untransfected neuron in the vicinity of neurons positive for LGI1-shRNA (Scale bar = 50 µm). **B)** Current injection protocol and AP shape. **C)** Excitability thresholds of neurons close to cells positive for LGI1-shRNA infection. Representative traces show the current injection from −10pA to 40pA for shRNA-LGI1 and scramble treatment. **D)** Bar graph showing the half-width of the first action potential at rheobase. **E)** Spike latency between the first four APs at 90pA. 90pA was chosen as the current step in which all the cells in the data base where spiking at least 4 times. **F, G)** Bar graph showing membrane input resistance and cell capacitance in of neurons close to cells positive for LGI1-shRNA and Scr-shRNA. Scr-shRNA: n = 12, LGI1-shRNA: n= 13 in 3 mice for each group.

### LGI1 downregulation affects circuits network excitability

Thus, subacutely decreasing LGI1 expression has an impact on synaptic transmission and neuronal excitability. Since LGI1 is a secreted protein, it is challenging to study the effect of a spatially restricted knockdown on broader network excitability within the hippocampus. We, therefore, took advantage of the more diffuse knockdown obtainable in primary hippocampal neuronal cultures. It has previously been shown that epileptic bursts and action potentials correlate with calcium transients in cultures (Kovac *et al.*, 2014). Therefore, we performed live fluorescence measurement of calcium activity in high-density cortical primary cultures (Figure 5 A). We showed that neurons treated with LGI1-shRNA were spontaneously more active than the control group (Scr-shRNA mean percentage of active neurons = 43.08 ± 3.87 %; LGI1-shRNA mean = 58.46 ± 5.16 %; p= 0.02, two-tailed Student’s t-test; Figure 5 C). There was also a trend for a higher spike frequency in the LGI1-shRNA group (Scr-shRNA mean = 0.08 ± 0.009; LGI1-shRNA mean = 0.1 ± 0.01; p=0.16, two-tailed Student’s t-test) (Figure 5 D).

**Figure 5:**
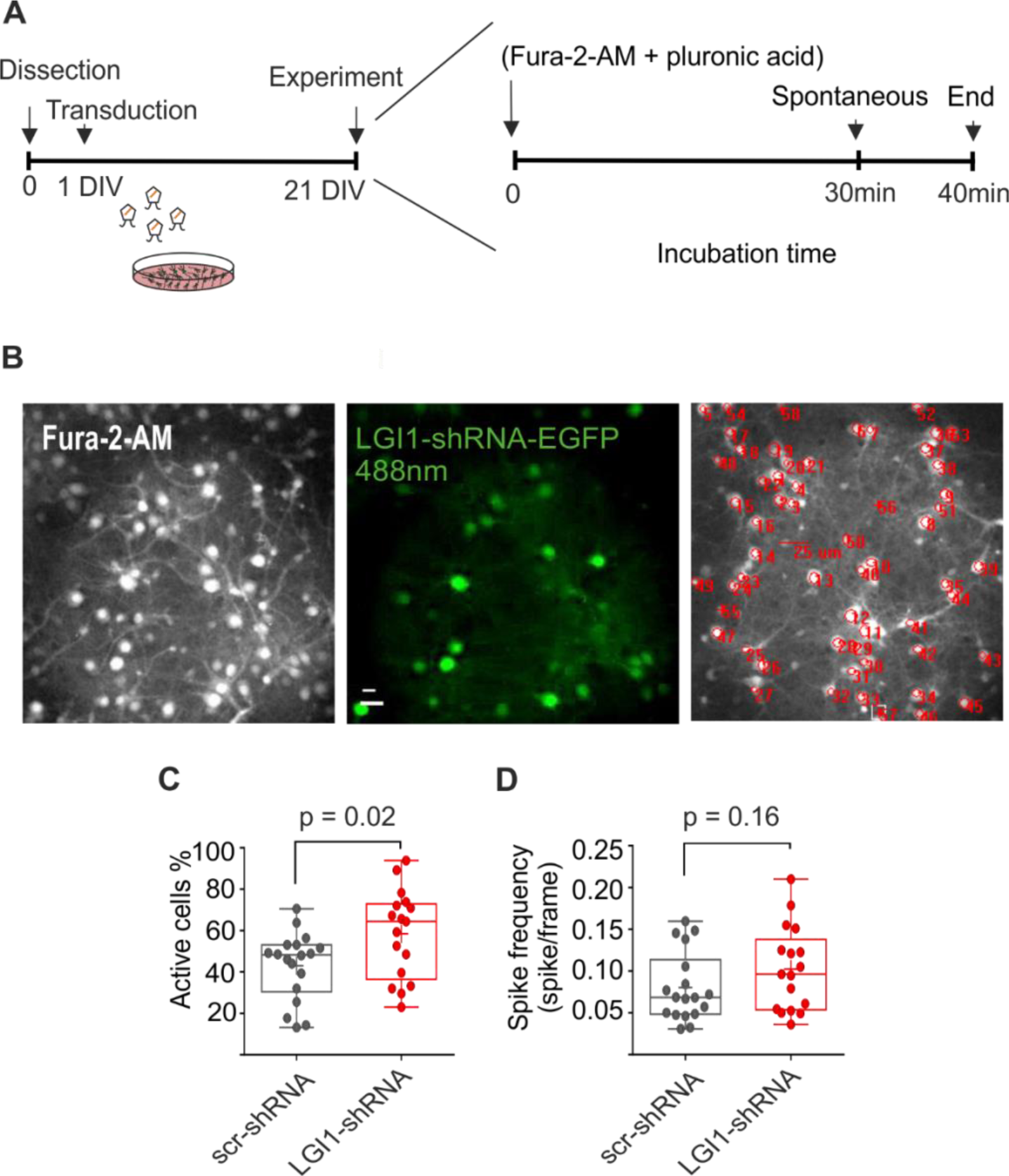
LGI1-shRNA affects basal calcium concentration in primary cultures. **A)** Protocol timeline for the live calcium imaging experiment. **B)** Representative 20X field of view of neuronal cultures treated with Fura-2-AM and pluronic acid (510 nm emission filter). The second picture shows the 488nm EGFP+ cells (scale bar = 25μm). The bottom picture is a representative snapshot of cells capturing Fura-2-AM within their cellular membrane. Red circles are manually selected ROI of their soma. **C)** Percentage of active cells in basal conditions **D)** Spike frequency for Scr-shRNA and LGI1-shRNA treatment in basal activity (Scr-shRNA: n = 18, LGI1-shRNA: n= 17 from 4 independent preparations).

Furthermore, we measured neuronal excitability using a multielectrode array (Figure 6 A, B). Hippocampal neuronal cultures were transduced at 1DIV and unit activity was recorded at 21DIV (Figure 6 A). We analysed the mean bursting rate (MBR), which was significantly higher in neurons transduced with LGI1-shRNA compared to the neurons transduced with Scr-shRNA (Scr-shRNA MBR mean = 5.61 ± 0.68 bursts/minute, LGI1-shRNA MBD mean = 8.92 ± 1.18 burst/minute, p = 0.04, two-tailed Student’s t-test) (Figure 6 C). Despite similar mean burst duration (MBD), spike frequency intra burst (MFIB) was higher in LGI1-shRNA transduced cultures than that in Scr-shRNA transduced cultures (Figure 6 D, E) (Scr-shRNA MBD mean = 135.4 ± 7.12 ms, LGI1-shRNA MBD mean = 132 ± 6.5 ms, p = 0.73, two-tailed Student’s t-test; Scr-shRNA MFR median = 91.3 spikes/second, Rank = 115.15, IQR = 19.42 spikes/second, LGI1-shRNA MFR median = 107.1 spikes/second, Rank = 93.08 spikes/second, IQR = 32.41 spikes/second; p = 0.03, two-tailed Mann-Whitney test). Further analysis showed that there was a tendency for the average MFR (mean firing rate) of Scr-shRNA transduced cultures to be less than that of LGI1-shRNA transduced cultures, although this did not reach significance (Scr-shRNA MFR median = 1.06 spikes/second, Rank = 3.86 spikes/second, IQR = 1.05, LGI1-shRNA MFR median = 1.79 spikes/second, Rank =6.23 spikes/second, IQR = 2.01; p = 0.07, two-tailed Mann-Whitney tests, Scr-shRNA: n= 18 wells, LGI1-shRNA: 22 wells).

**Figure 6.**
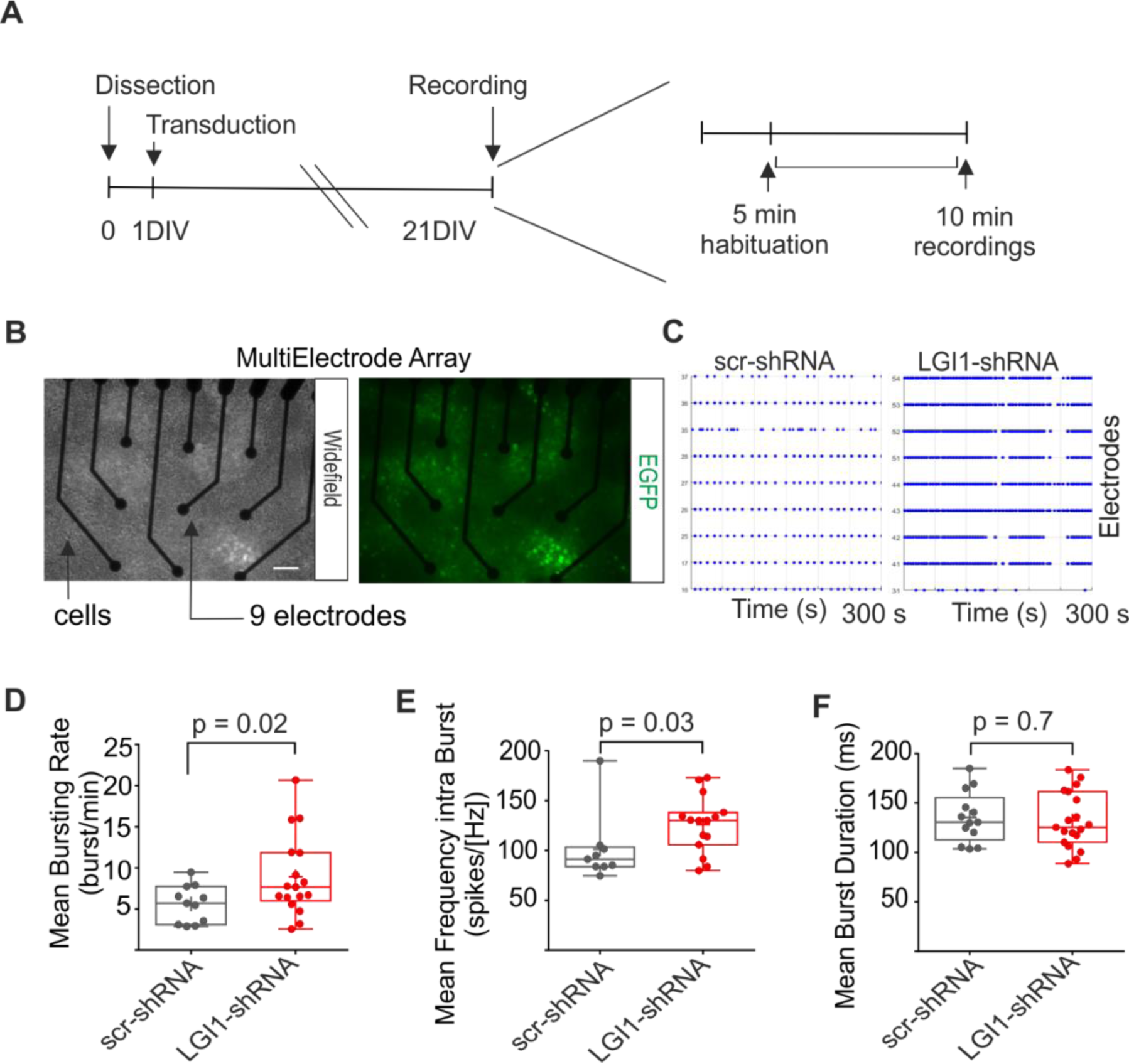
Subacute knock down of LGI1 affects network burst activity. **A)** Timeline of the experiments **B)** Representative picture of a single well of a multielectrode array. The 9 recording electrodes are in black, over a field of neuronal networks. Each electrode is of 30μm diameter. The picture shows 21 days old MEAs (left: wide field, right in green: expression of EGFP in transduced cultures, bottom: merged picture. Scale bar = 60μm) and **C)** raster plot (spikes) of the neuronal activity detected by single electrodes in LGI1-shRNA and Scr-shRNA. **D)** Mean bursting rate (Scr-shRNA: n= 11 wells, LGI1-shRNA: 17 wells; p = 0.04, Student’s t-test). **E)** Mean firing intra burst (Scr-shRNA: n= 11 wells, LGI1-shRNA: 17 wells; p= 0.03 Mann-Whitney test). **F)** Mean bursting duration (Scr-shRNA: n= 11 wells, LGI1-shRNA: 17 wells; p = 0.73 Student’s t-test). Data were plotted and averaged from 5 independent preparations.

### LGI1 downregulation does not affect cellular viability and number of excitatory synapses *in vitro*

Since LGI1 has been reported to be important for glutamatergic synapse development, we investigated the number of active excitatory synapses of *in vitro* hippocampal neurons. Co-localization of vGLUT1 (Vesicular Glutamate Transporter 1), a presynaptic marker of excitatory synapses, and Homer1, a post synaptic marker of active excitatory synapses, did not reveal any significant difference between neurons transduced with Scr-shRNA or LGI1-shRNA (Mander’s coefficient Scr-shRNA = 0.6 ± 0.04; LGI1-shRNA = 0.62 ± 0.03; p = 0.58 two-tailed t-test; Scr-shRNA: n = 11, LGI1-shRNA: n = 12) (Supplementary figure 4 A, B). Neuronal death in cultures and brain slices is an important mechanism of homeostasis (Fricker *et al.*, 2018). To check whether removal of LGI1 in cultures affects cell viability which could lead to the difference in calcium signaling and burst activity, we used a mortality cell assay with propidium iodide staining (PI) (Ayşit-Altuncu *et al.*, 2018). The mortality rate between LGI1-shRNA transduced and Scr-shRNA transduced cultures were comparable (Scr-shRNA mortality rate median = 51.38%, Range = 44.9% (38 to 82.9%), IQR = 10.68 %, LGI1-shRNA mortality rate median = 48.25%, Range = 25.3% (42.1 to 6.4%), IQR = 12.38%; p= 0.9, Mann-Whitney two-tailed test) (Supplementary Figure 4 C, D).

## Discussion

Extracellular matrix proteins have recently emerged as key contributors to the maintenance and regulation of physiological processes (Ferrer-Ferrer and Dityatev, 2018). Here, we investigated the role of LGI1, a trans-synaptic secreted protein, part of the synaptic ECM, after its subacute depletion in neuronal networks

In this work, we established a RNA silencing method to downregulate LGI1 subacutely in both *ex vivo* hippocampal slices and neuronal culture. We demonstrated that subacute removal of LGI1 in MF-CA3 hippocampal circuits was able to alter local network and neuronal excitability. Pharmacological experiments with a blocker of Kv1 ion channels indicated that modification of Kv1 was the likely basis of the observed increased facilitation observed in MF-CA3 circuits. Moreover, we demonstrated that a 40% downregulation of LGI1 in primary neuronal cultures was sufficient to alter the basal calcium levels and network bursts, without affecting neuronal survival.

So far, investigation of LGI1 in the physiology of neuronal circuits has been explored with the use of transgenic animals or with acute applications of auto-antibodies against LGI1 in primary cultures and slices (Fukata et al., 2016). These models demonstrated that LGI1 plays a crucial role during embryonic development and modulates both pre and post synaptic transmission. Several discrepancies in the effects described from multiple groups could be ascribed to the different genetic background of the animal models used, the differing ages of the brain samples examined and the diverse anatomical zones investigated (Chabrol et al., 2010a; Fukata et al., 2010a; Yu et al., 2010). Moreover, some LGI1 mutations cause a loss of functional monomer, while others have a dominant-negative effect, hence triggering epilepsy probably via two different mechanisms of action. Finally, experiments with auto-antibodies from patients with LGI1 limbic encephalitis suggest that acute interference with LGI1 is sufficient to alter neuronal circuits and synaptic transmission, without a significant immune/inflammatory component (Lalic et al., 2011; Ohkawa et al., 2013; Petit-Pedrol et al., 2018).

In order to separate the contribution of developmental issues and of the inflammatory response from the removal of LGI1 protein in neuronal circuits, we designed and validated a subacute method based on small interfering RNA for downregulation of LGI1, avoiding the use of mutants or of polyclonal IgGs from encephalitic patients. To our knowledge, this approach has not been used before in the investigation of the role of LGI1 but it can establish the effects of subacute LGI1 downregulation. We achieved ~40% knockdown of the endogenous LGI1 protein (Fukata et al., 2006; Kunapuli et al., 2009; Zhou et al., 2018). This knockdown level is close to a heterozygous state, helping to dissect realistic physiological consequences of LGI1 molecular alterations on neuronal circuits.

Despite evidence that LGI1 has a pivotal role in glutamatergic neurons, LGI1 staining is also present in interneurons and glia (Fukata et al., 2016). Therefore, in this study, we used a broad promoter (mU6) for silencing RNA activity which is able to induce a constant production of the shRNA in any cell type.

Measurements of short-term plasticity using *ex vivo* hippocampal slices previously injected with LGI1-shRNA, showed greater facilitation of MF to CA3 transmission. This result is crucial to attribute functional alterations of excitatory transmission to subacute downregulation of LGI1, without the contribution of developmental circuits abnormalities or temporal lobe inflammation. These results are in line with previous work that measured higher spontaneous burst activity and interictal-like depolarisation by extracellular field recordings in CA1 of LGI-KO (Yu et al., 2010; Boillot et al., 2016; Petit-Pedrol et al., 2018). Also, the acute application of VGKC-IgGs in the DG-CA3 pathway confirmed that acute interference with LGI1 provoked higher neurotransmitter release from the granule cells and increased postsynaptic excitability (Lalic et al., 2011). The effects that we observed on low-frequency facilitation at the mossy fiber-CA3 synapse suggest that initial activation of presynaptic Kv1.1 during low-frequency facilitation restricts the degree of facilitation. Increasing the inactivation of Kv1.1 by removing LGI1 thus removes this “brake” and so enables greater low-frequency facilitation. Loss or a reduction in LGI1 could thus act as an initiator of seizures and could facilitate propagation in the hippocampus, as previously hypothesized (Seagar et al., 2017). The fact that an extracellular structural protein influences network excitability and glutamate release has been debated for many years. The first evidence that LGI1 indirectly has a role in cellular excitability arrived when it was first co-purified in the macromolecular structure formed by Kv1.1 presynaptic potassium channel and postsynaptic AMPAR. Since then, it has been suggested that LGI1 influences Kv1.1 channels by two mechanisms: a prolongation of its opening state by interference with the Kvβ1 regulatory subunit, and an influence on the number of Kv1.1 channels recruited to the presynapse. Inside-out patches in oocytes showed that mutations or removal of LGI1 proteins lead to a shortening of the opening time of Kv1.1 channels. This has been attributed to the ability of a single kvβ1 unit to block the whole Kv1.1 tetramer. Instead, multiple LGI1 are required to interfere with kvβ1, so that a small decrease in LGI1 relative to kvβ1 causes a much greater effect than expected (Schulte et al., 2006). *Ex vivo* LGI1 reduction affected short-term plasticity of MF to CA3 synapses, resulting in a significant over facilitation during repetitive stimulation. This is probably caused by early inactivation of Kv1.1 channels since blocking Kv1 with α-DTX occluded this effect. Therefore, it is also possible that subacute reduction of LGI1 proteins causes alteration of neurotransmission by functional modification of Kv1.1 kinetics. We cannot, however, completely exclude a post-synaptic contribution from LGI1 reduction. Application of α-DTX in both Scr-shRNA and LV-shRNA, caused a small increase in the amplitude of the response (in slope). At the same time, we measured also a trend to a decrease in the fiber volley slope, implying fewer axons recruited by the stimulation. Together these results indicate that the release probability for any single fiber may be increased, suggesting that Kv1.1 could be contributing to the initial release probability. Our findings can, therefore, be explained by the fact that the reduction of LGI1 promotes inactivation of Kv1.1 during the wind-up resulting in a greater release probability.

Patch-clamp experiments of cells near to EGFP+ neurons in LGI1-shRNA slices revealed that reduction of LGI1 affects the physiological properties of neurons close by transduced LGI1-shRNA positive cells. Rescue experiments in previous work supported that LGI1 probably has paracrine effects onto neighbouring neurons, hence local and circumscribed reduction may affect other neurons too (Lovero *et al.*, 2015). Our data show that subacute reduction of LGI1 reduces AP threshold (Seagar *et al.*, 2017). This was also observed with acute incubation of VGKC-IgG by another group (Lalic *et al.*, 2011). LGI1 subacute reduction also decreases the inter spike interval, but does not affect the AP half-width. This is probably due to the expression pattern of Kv1.1, which is more abundant at the AIS and axons (Robbins and Tempel, 2012). We asked if these changes in excitability that we observed translated to an effect on network excitability. We employed calcium measurements and MEA devices to show, for the first time, that subacute LGI1 reduction results in an alteration in neuronal network excitability. The measurement of calcium waves is of critical importance since they are strongly correlated with epileptic-like bursts and action potentials *in vitro* (Kovac et al., 2014). These initial studies indicated that broad heterozygous-like LGI1 downregulation was sufficient to alter basal calcium levels and burst activity of *in vitro* neuronal networks. These results corroborate previous findings from studies of LGI1 mutants in cell lines (Nobile et al., 2009; Pakozdy et al., 2014) and extend the present knowledge about LGI1-IgGs. Interestingly, only one previous study investigated calcium waves in neuronal cultures showing the opposite effect to what we showed here. This might be attributable to long term (24 -72 hours) IgG incubations leading to neuronal toxicity in that study (Ayşit-Altuncu et al., 2018). To conclude, the results reported here demonstrate that a subacute decrease of LGI1 can result in alterations of the DG-CA3 pathway excitability and increases in network hyperexcitability *in vitro*. Our findings support new pathophysiological mechanisms by which alteration of LGI1 affects brain excitability and opens up new clinical intervention routes through, for example, the use of small molecules that increase Kv1.1 inactivation (Niespodziany et al., 2019).

## Acknowledgments

This work was supported by Marie Skłodowska-Curie grant agreement 642881 (MW, AD and EL), the Medical Research Council (MW), and the Marie Skłodowska-Curie Individual Fellowships and Epilepsy Research UK (GL). We would like to thanks all the Kullmann Lab for the helpful discussions, Dr Erica Tagliatti for the help with neuronal culture and imaging, Dr Jenna Carpenter for her advice with the molecular strategy, Dr Tawfeeq Shekh-Ahmad for his help with calcium imaging, Dr Gareth Morris, Dr Marion Mercier, Dr Vincent Magloire and Dr Sylvain Rama for their help with electrophysiology and Dr Antonia Yam and Dr Ilaria Colombi for their help with multi-electrode arrays optimization and analysis.

---

**Supplementary figure 1.**
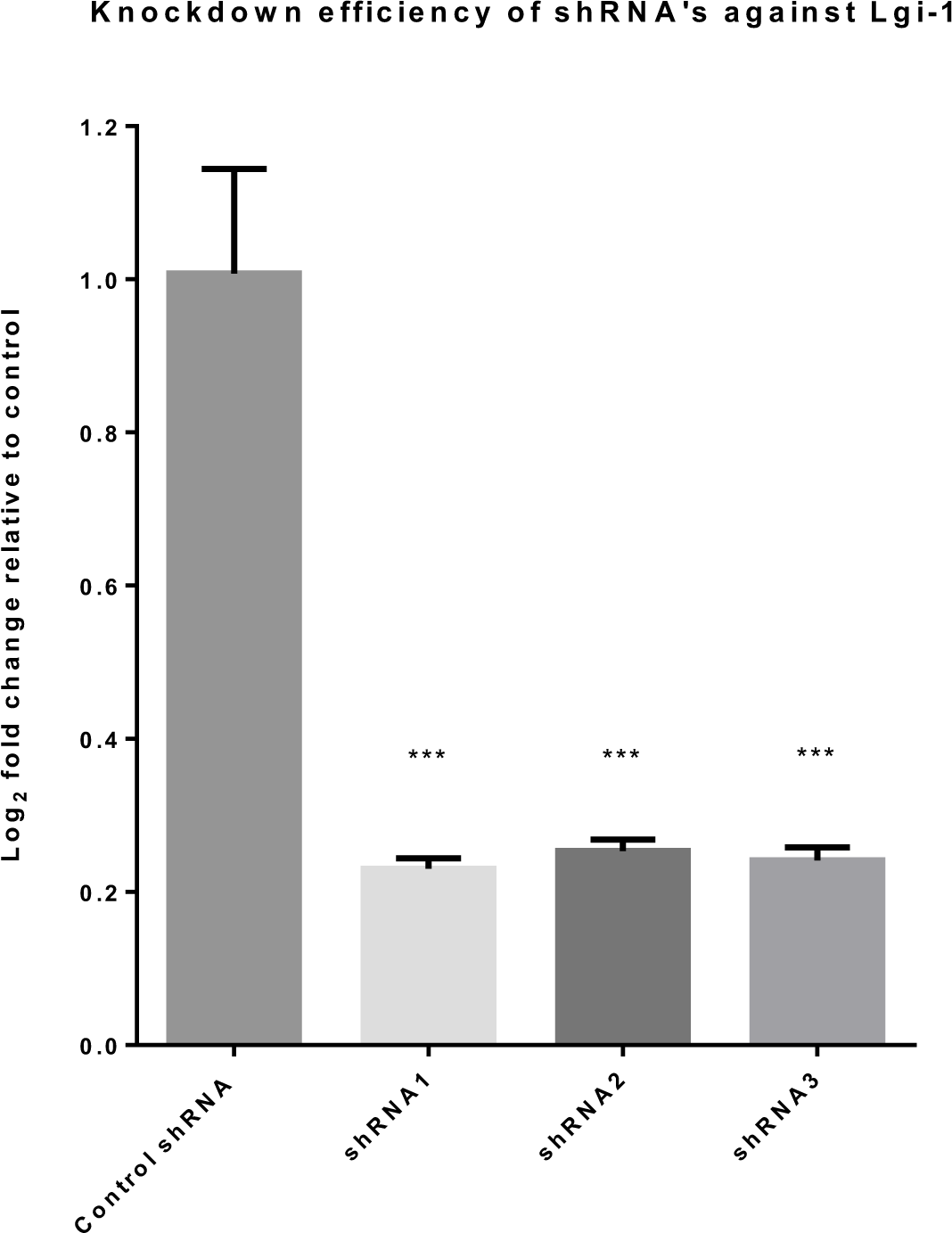
shRNA mediated knockdown of Lgi-1 in N2a cells: All 3 shRNA plasmids after 48 hours of transfection were able to significantly knockdown mouse Lgi1 in mouse neuroblastoma cell line Neuro2a (One-way ANOVA with Bonferroni correction for comparisons F (3, 20) = 174.0, n=3)

**Supplementary figure 2.**
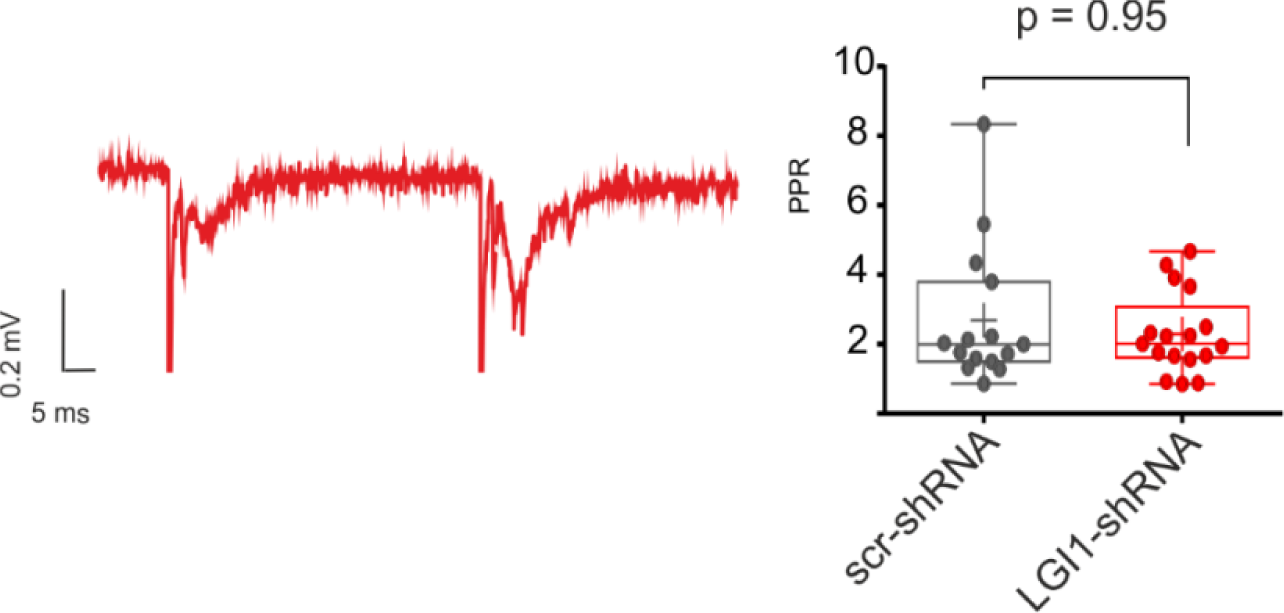
LFP Paired pulse ratio. *Representative traces with a bar graph illustrating no difference in paired-pulse ratio between groups.* Scr-shRNA: n = 15 from 5 mice and LGI1-shRNA: n= 17 from 8 mice.

**Supplementary figure 3.**
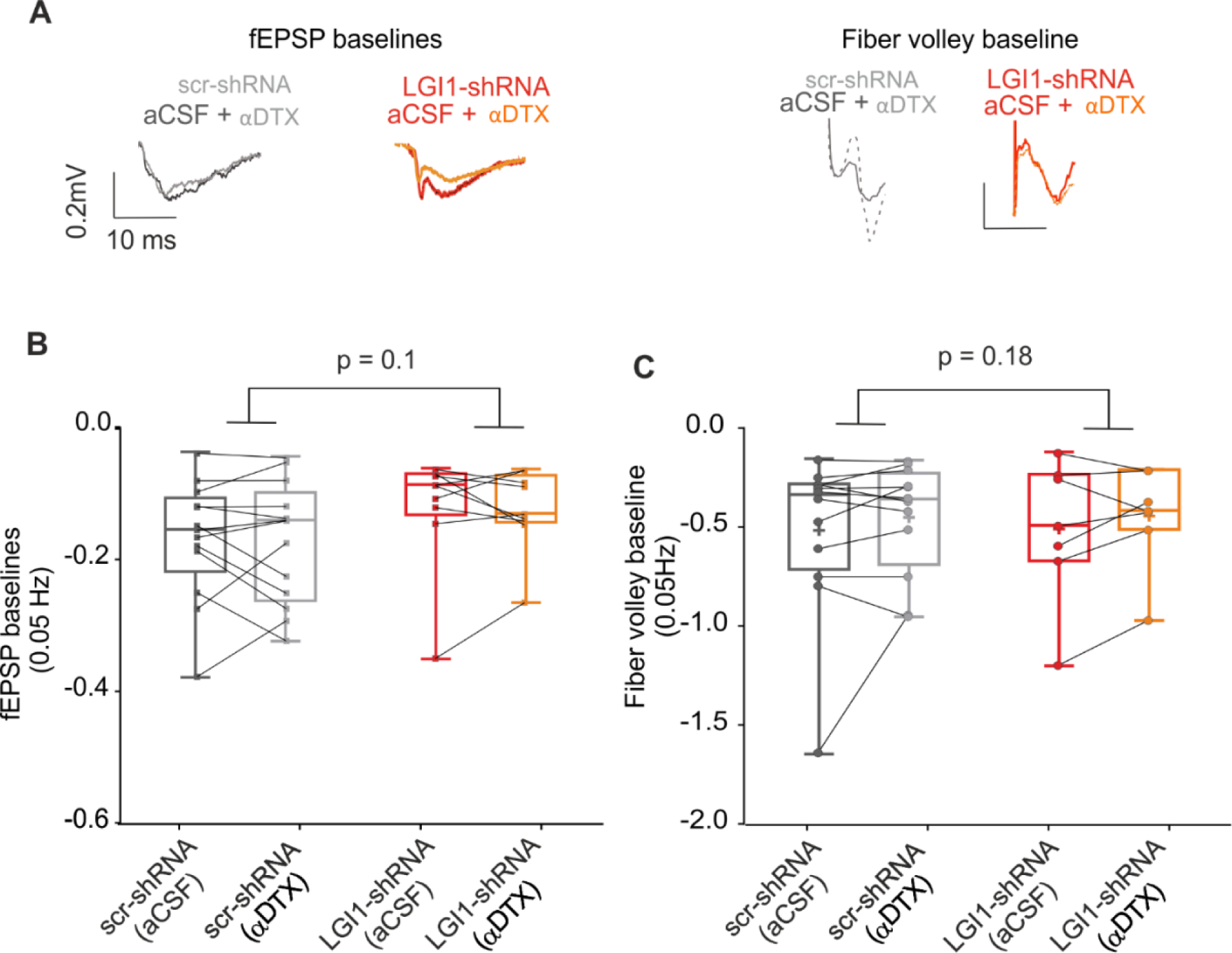
Baseline and fiber volley are not significantly different before and after the application of alpha dtx. **A)** scatter plot illustrating the slope of the amplitude relative to the baseline stimuli across groups. **B)** scatter plot illustrating the fiber volley slope during baseline stimulations (0.05Hz) across groups.

**Supplementary figure 4.**
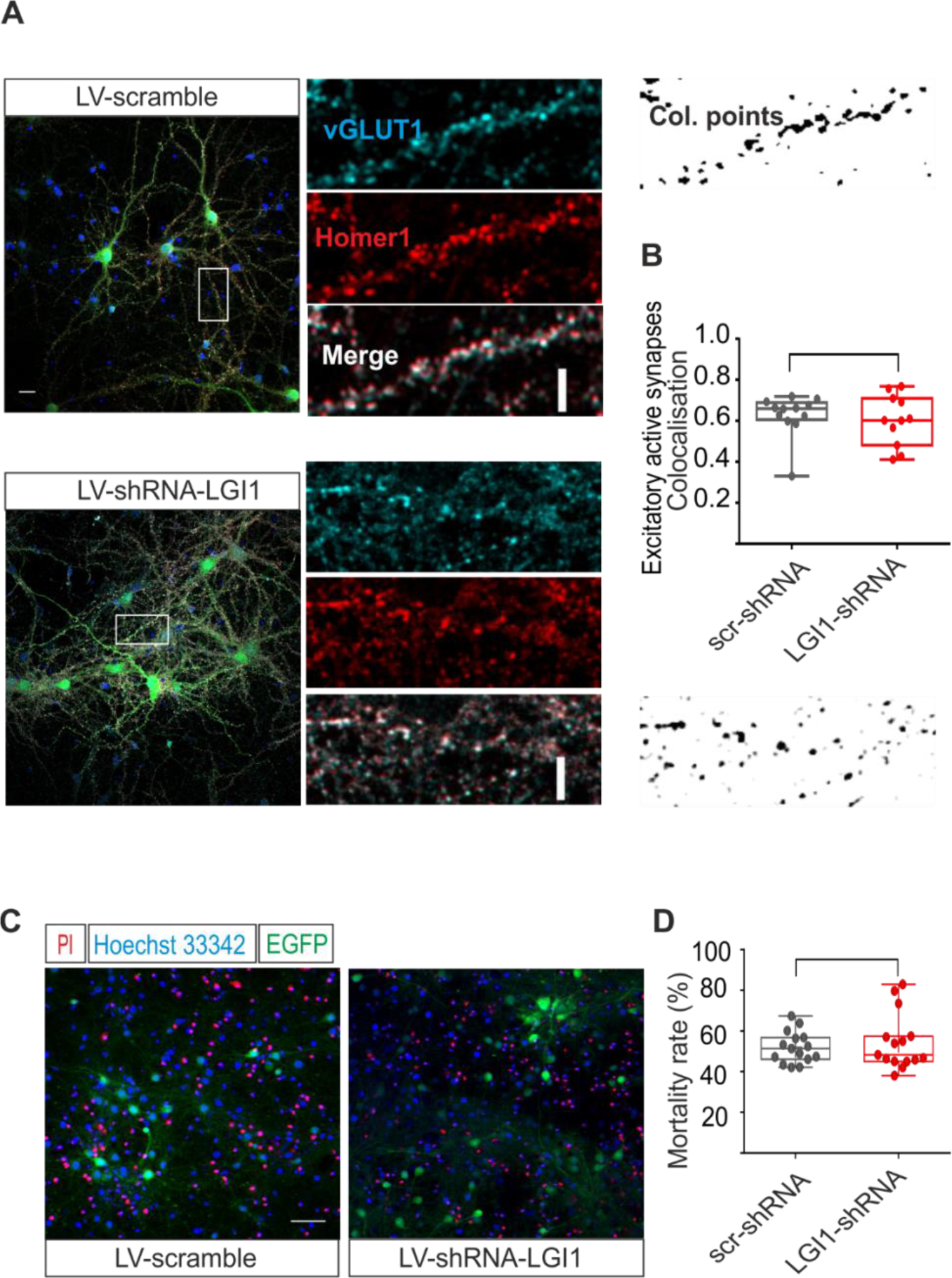
Acute knock down of LGI1 does not affect either cellular viability nor the number of active excitatory synapses in primary cultures. **A)** Representative image of neuronal cultures and zoom of a dendrite stained for excitatory synapses and **B)** quantification of excitatory synapses colocalisation. Green = LGI1-shRNA or Scr-shRNA transduced positive neurons, blue = Hoechst 33342 nucleus staining; cyan = vGLUT1 presynaptic excitatory marker, red = Homer1 post synaptic marker active excitatory synapses. Scale bar is 20 μm. *(Mander’s coefficient Scr-shRNA = 0.60 ± 0.037; LGI1-shRNA = 0.62 ± 0.029; p = 0.584, two-tailed t-test; Scr-shRNA: n = 11, LGI1-shRNA: n = 12)* **C)** representative image of propidium iodide staining in primary cultures. Red particles are the nuclei of dead cells with the damaged cellular membranes, blue represents the nuclei of all the cells in the coverslips and green neurons are those transduced either by Scr-shRNA or LGI1-shRNA virus (scale bar = 50 μm). **D)** The quantitative plot of mortality rate (red cells / blue cells) between Scr-shRNA and LGI1-shRNA treated cultures at DIV21. The experiments were performed in triplicates for a total of 3 different preparations. (scr-shRNA n = 9 coverslips, LGI1-shRNA n= 9 coverslips from three independent preparations)

**Supplementary table 1:**
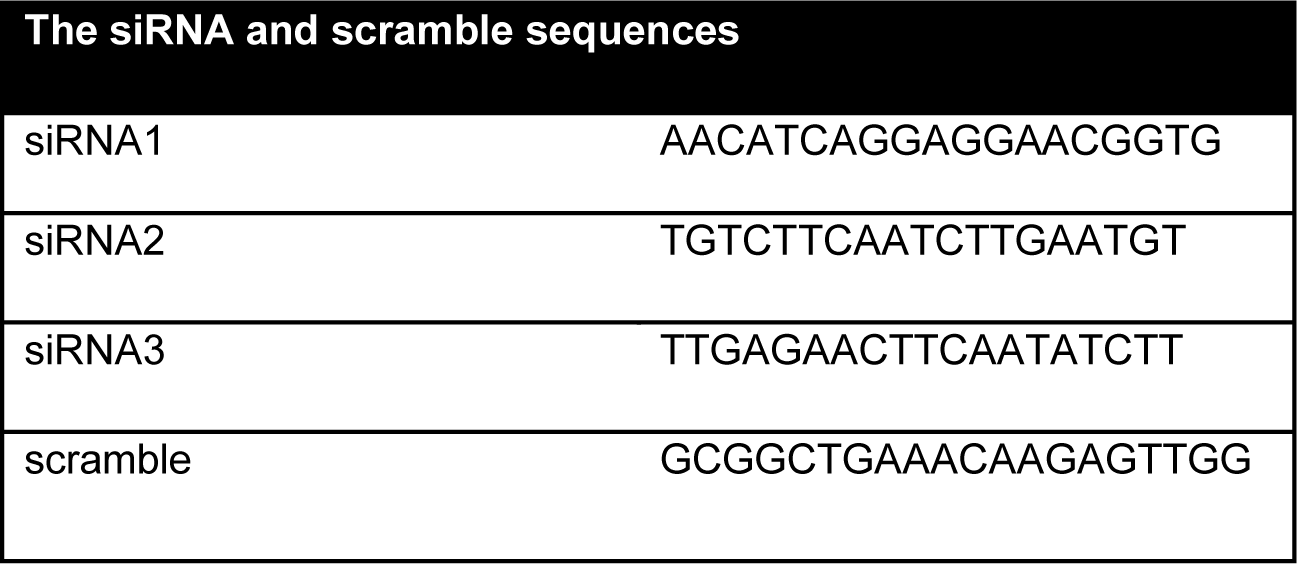
siRNA for LGI1 and scramble sequences

